# Riemannian Geometry for the classification of brain states with fNIRS

**DOI:** 10.1101/2024.09.06.611347

**Authors:** Tim Näher, Lisa Bastian, Anna Vorreuther, Pascal Fries, Rainer Goebel, Bettina Sorger

## Abstract

**Background:** Functional near-infrared spectroscopy (fNIRS) has recently gained momentum as a reliable and accurate tool for assessing brain states. This increase in popularity is due to its robustness to movement, non-invasive nature, portability, and user-friendly application. However, compared to functional magnetic resonance imaging (fMRI), fNIRS is less sensitive to deeper brain activity and offers less coverage. Additionally, due to fewer advancements in method development, the performance of fNIRS-based brain-state classification still lags behind more prevalent methods like fMRI.

**Methods:** We introduce a novel classification approach grounded in Riemannian geometry for the classification of kernel matrices, leveraging the temporal and spatial channel relationships and inherent duality of fNIRS signals—more specifically, oxygenated and deoxygenated hemoglobin. For the Riemannian geometry-based models, we compared different kernel matrix estimators and two classifiers: Riemannian Support Vector Classifier and Tangent Space Logistic Regression. These were benchmarked against four models employing traditional feature extraction methods. Our approach was tested in two brain-state classification scenarios based on the same fNIRS dataset: an 8-choice classification, which includes seven established plus an individually selected imagery task, and a 2-choice classification of all possible 28 2-task combinations.

**Results:** The novel approach achieved a mean 8-choice classification accuracy of 65%, significantly surpassing the mean accuracy of 42% obtained with traditional methods. Additionally, the best-performing model achieved an average accuracy of 96% for 2-choice classification across all possible 28 task combinations, compared to 78% with traditional models.

**Conclusion:** To our knowledge, we are the first to demonstrate that the proposed Riemannian geometry-based classification approach is both powerful and viable for fNIRS data, considerably increasing the accuracy in binary and multi-class classification of brain activation patterns.

## **1** Introduction

Functional near-infrared spectroscopy (fNIRS) is an emerging functional neuroimaging technique that has recently gained momentum due to its non-invasive nature, portability, silent data acquisition, robustness against movement artifacts, and a favorable balance between spatial and temporal resolution.^1–3^ FNIRS detects hemodynamic responses, which are also measured by functional magnetic resonance imaging (fMRI) and have there been linked to changes in neuronal activity.^4^

Both fMRI and fNIRS assess neuronal activity indirectly via the blood-oxygen-level-dependent (BOLD) signal. The BOLD signal is especially linked to neuronal gamma oscillations.^5, 6^ Intriguingly, the BOLD signal reflects changes in both gamma and neuronal firing rates when they are driven by external stimuli; yet when changes are due to internal state fluctuations, they are captured by BOLD specifically for gamma and much less for firing rates.^7^ This makes gamma oscillations a crucial marker for studying brain activity, particularly in the context of internal state changes.^5, 8^ Unlike fMRI, fNIRS measures two distinct hemoglobin signals: oxygenated (HbO) and deoxygenated (HbR) hemoglobin. Although these signals are naturally anticorrelated,^9^ they are not merely redundant; rather, they can provide complementary information about brain activation.^10^ Despite measuring two separate signals, the accuracy of fNIRS-based classification of brain states has yet to reach the level of fMRI. This disparity could also be attributed to the fact that fMRI benefits from more advanced and well-developed data analysis techniques due to its more frequent use compared to fNIRS. Consequently, it is likely that the full potential of fNIRS data analysis has not yet been realized, given its historically less developed and less widely known status. While fMRI can classify brain states by utilizing spatial differentiation of multivariate activation patterns across the whole brain,^11^ fNIRS is constrained by its limited spatial resolution and coverage, particularly limited in depth penetration. As a result, not all classification methods suited for fMRI data may perform as effectively when applied to fNIRS data.

The current classification methods for fNIRS-based brain-state identification are often adapted from other functional neuroimaging techniques but are not specifically tailored to the unique properties of fNIRS signals. Traditional classifier approaches include Linear Discriminant Analysis (LDA), Support Vector Classifier (SVC), Random Forest (RF), and Logistic Regression (LR), which typically rely on channel-based features like mean activation, channel variance, regression slope, and zero crossings.^12, 13^ While more advanced methods, such as convolutional and long-/short-term memory (LSTM) neural networks, have been employed to improve classification accuracy, these deep-learning approaches require large amounts of training data, which are often not available.^14^

Given these limitations, there is a need to develop methods specifically tailored to the unique properties of fNIRS. In practice, this involves emphasizing the dual nature of fNIRS signals by utilizing the potentially complementary information in HbO and HbR data. Increased specialization could bridge the gap between the practicality of fNIRS and the precision seen in fMRI, making fNIRS a more viable alternative for brain-state classification. To achieve this, we combined HbO and HbR signals in block diagonal matrices and subsequently applied Riemannian Geometry-based classification models. Instead of relying on features from individual channels, Riemannian Geometry-based models utilize relationships between channels. To our knowledge, we are the first to demonstrate that the classification of fNIRS data using Riemannian Geometry offers a promising alternative to other machine-learning methods. The framework offers well-established tools that allow for the independent manipulation and therefore tuning of separate signals within the same model. This allows us to effectively capture spatial co-activation patterns separately for both HbO and HbR with only one model. Additionally, we recorded participants’ responses regarding the ease and comfort of each task, as more favorable tasks could be generally better suited for classification. Our approach outperforms traditional methods in both 8-choice and 2-choice classification scenarios. While Riemannian Geometry is a well-established technique in other classification fields,^15–21^ we show that it is also a powerful and viable option for fNIRS data.

## 2 Method

### 2.1 Optode setup and General Procedure

#### 2.1.1 Participants

Seven healthy participants (mean age: 32.1 +/- 10.6 years, three female) were included in this experiment (Supplementary Table 1). All participants were either externally recruited or students and staff members of the Faculty of Psychology and Neuroscience at Maastricht University. Written informed consent was acquired before the experiment. The local ethics committee approved the experiment (Ethics Review Committee Psychology and Neuroscience). Upon successful completion of the experiment, participants received financial compensation.

#### 2.1.2 General Procedure

Before the experiment, participants received instructions via email (see Supplementary Table 2), including short but precise mental-task descriptions to train the task performance beforehand. Participants were asked to familiarize themselves with these information and devise their “own (mental) paradigm” (OP): A suitable mental task or process that was not included in the provided task set. The individual task should be familiar to them and they should feel comfortable executing it upon being cued. During the fNIRS session, each participant was seated in a darkened and sound-attenuated room. The fNIRS cap was placed on the participant’s head, and all optodes were adjusted to ensure orthogonality to the scalp and stable optode-scalp contact. During the optode placement, participants were presented with the same stimulation (task cues) used for the experiment to practice the mental task performance until they felt comfortable. Participants were also instructed to get into a comfortable position and to relax their jaw and facial muscles during the recording session. The fNIRS session lasted approximately 1.5 h including participant preparation and data acquisition runs. At the end of the session, participants rated the pleasantness and easiness of all eight mental task performances on a 10-point Likert scale (i.e., 1 = extremely unpleasant/extremely difficult to 10 = extremely pleasant/extremely easy).

#### 2.1.3 Experimental Runs

Participants performed four runs with a total of eight mental imagery tasks (Fig 1b). Mental imagery tasks included Mental Talking (MT), Spatial-Navigation Imagery (SN), Mental Drawing (MD), Mental Singing (MS), Mental Calculation (MC), Mental Rotation (MR), Tennis Imagery (TI), and the previously devised Own Paradigm (OP) (Supplementary Table 2). Participants were instructed to engage in the mental task cued on a screen for the duration that the cue was shown. To instruct participants when to engage in mental imagery, three letters of the mental task name were displayed (e.g., ”dra” instead of ”mental drawing”; other abbreviations in Supplementary Table 2). This was meant to keep visual input similar across tasks and avoid visual stimulation-based differences in cerebral activation. Instructions were presented for 2s, succeeded by a fixation cross. Participants were told to engage in the mental task displayed on the screen with the greatest possible consistency and to stop when the instruction to rest (“res”) was given. During 20s rest, participants were asked to refrain from engaging in any specific thoughts or tasks. Each imagery task was performed for a total of twelve trials (trial duration: 25s-35s; encoding duration: 5s-15s), resulting in 96 trials per participant (Fig 1b). For each participant, each run contained two conditions that were performed in an alternating order (Fig 1b). Pairings were pseudo-randomized to avoid the same pairings across participants.

**Fig 1.**
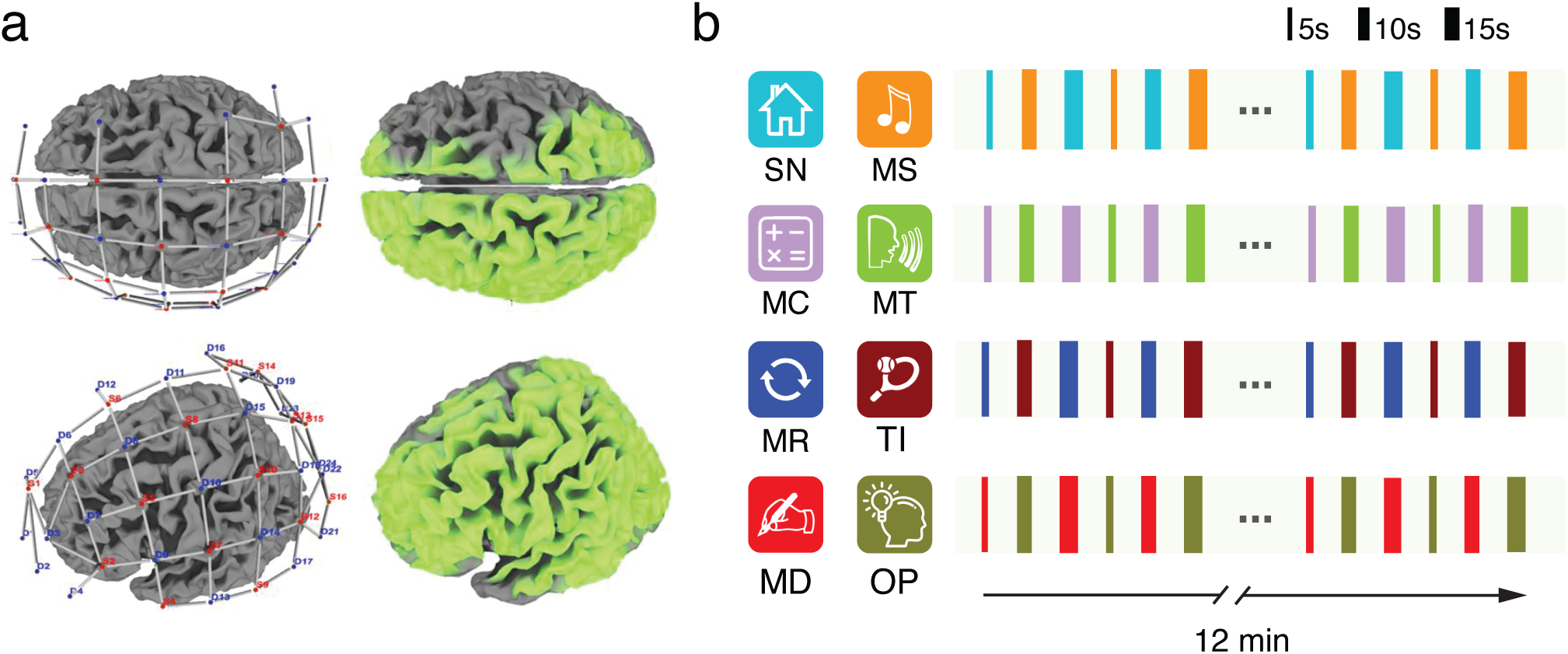
Optode setup with resulting light-intensity map and experimental procedure. **a** Optode setup (left) and light-intensity map of the montage (right) that indicates the coverage of the fNIRS sensors (light green regions). **b** Example structure of task pairs used in the four functional runs for one participant. The task pairings varied across participants. Each run started with a one-minute rest period. Task pairs were presented in alternation for 24 trials with 20-s rest periods after each task trial. Trial durations varied between 5s, 10s, and 15s. This pattern was repeated four times in each run and resulted in a total duration of 12 min per run. Abbreviations: MC: mental calculation, MD: mental drawing, MR: mental rotation, MS: mental singing, MT: mental talking, OP: own paradigm, SN: spatial-navigation imagery, TI: tennis imagery

#### 2.1.4 Data Acquisition

Data were acquired with a NIRScout-816 system (NIRx Medizintechnik GmbH, Berlin, Germany) with 16 source and 24 detector optodes (Fig 1a). Detectors and sources were positioned according to the international 10-20 Electroencephalography (EEG) system. The following list contains the optode positions according to the 10-20 system. Source positions were over AFz, F5, FT7, FC3, FCz, C5, C1, TP7, CP3, CPz, P5, P1, P2, POz and O1. Detectors were positioned over FPz, AF3, AF7, AF4, Fz, F3, FC1, FC5, C3, Cz, FC2, T7, CP5, CP1, CP2, P7, P3, Pz, P4, PO7, PO3, PO4, and Oz. The optodes covered the whole left hemisphere and potential task-relevant brain regions within the right hemisphere. This montage was chosen to include areas where we expected to observe imagery task-related brain activity. Data were acquired using NIRStar 15.2 (NIRx Medizintechnik GmbH, Berlin, Germany) at 3.47 Hz.

### 2.2 Data Analysis and Models

#### 2.2.1 Data Preprocessing

All preprocessing steps were executed using Satori (v2.0). Data were trimmed to 5s before the onset of the first mental task trial and 15s after the offset of the last trial. The raw time course values were converted to HbO and HbR concentration values using the modified Beer-Lambert law. According to the optode setup, the data were split in 62 channels for Hbo and 62 channels for HbR. Subsequent motion correction included spike removal (ten iterations, 5s lag, 3.5 threshold, 0.5 influence) with monotonic interpolation and Temporal Derivative Distribution Repair (TDDR), preserving high-frequency components. Due to hardware limitations, we did not obtain time courses from short channels, which is why we were not able to perform a short-separation regression to correct for systemic noise. Instead, a global component regression based on principle component analysis was performed, removing the first component. Finally, the signals were filtered using Gaussian low-pass (0.4 Hz) and Butterworth high-pass (0.01 Hz), and z-scored. Because of the higher sampling rate of fNIRS compared to fMRI, we expected to capture more detailed temporal dynamics of the hemodynamic response function. We therefore cut the data from cue onset to 10s post cue for classification to include the initial dip and the peak response. Within this 10s window, we estimated several features per channel, including mean, variance, kurtosis, and others. These features were then input into the traditional classifiers. Hyperparameters related to the classifiers underwent hyperparameter tuning to optimize their performance (Supplementary Table 3). For the Riemannian methods, different kernel matrices were estimated from the 62 HbO and HbR channels. The estimation of these kernel methods was according to several different covariance and (non)-linear kernel functions which were also considered hyperparameters of the Riemannian models. The resulting matrices served as features for the Riemannian classifiers, which also underwent hyperparameter tuning.

#### 2.2.2 Classifiers

We compared twelve classifiers: eight Riemann-based models and four traditional models. The traditional models were: LDA, LR, SVC, and RF. We computed standard features (mean, variance, peak, zero-crossings, skew, and kurtosis) for these models as described in the fNIRS literature.^22^ A full table of models and their hyperparameters can be found in Supplementary Table 3.

#### 2.2.3 Kernel matrix estimation

For EEG-based BCIs, classification based on covariance matrices is standard practice. These matrices, specifically termed ”channel covariance matrices,” are derived from the covariances between channel pairs. Covariance measures the extent to which two time series vary together: a positive covariance indicates that increases in one time series are matched by increases in the other, whereas a negative covariance suggests that an increase in one corresponds to a decrease in the other. However, a covariance of zero does not necessarily denote independence between the time series; rather, it indicates that there is no linear relationship between them.

Besides simple covariance, one can also compute more generalized measures. These measures, often realized through kernel functions, are essential in modern machine learning as they capture relationships in the data that extend beyond linear associations.^23^ Kernel functions enable the modeling of non-linear interactions by mapping the data to a higher-dimensional space. In this new space, the non-linear associations can be more effectively identified and therefore leveraged for classification tasks. Mathematically, a kernel matrix **K** is defined such that each element *K_ij_* measures the similarity or dependence between observations **x***_i_* and **x***_j_* according to a predefined function. In line with this definition, covariances between channels can be considered a type of kernel. Since we estimate both covariances and kernel functions, we collectively refer to all matrices representing relationships between channel data as ”kernel matrices.”

We used several different kernel functions to obtain the kernel matrices between channels, employing the following estimators: the correlation coefficient matrix (“corr”), the numpy-based covariance matrix (“cov”), Huber’s M-estimator based covariance matrix (“hub”), the Ledoit-Wolf shrunk covariance matrix (“lwf”), the oracle approximating shrunk covariance matrix (“oas”), the Schaefer-Strimmer shrunk covariance matrix (“sch”), the sample covariance matrix (“scm”), Tyler’s M-estimator based covariance matrix (“tyl”), the polynomial kernel (“poly”), which adds non-linearity by raising the dot product to a specified power, the radial basis function (RBF) kernel (“rbf”), which measures similarity based on the distance between input vectors; the Laplacian kernel (“laplacian”), which is similar to the RBF kernel but uses the L1 norm for distance measurement, and the cosine kernel (“cosine”), which measures the cosine of the angle between input vectors. All kernel functions were implemented using scikit-learn^24^ and pyriemann.^25^

Let **K**_HbO_ be the kernel matrix for the HbO channels, and **K**_HbR_ for the HbR channels, where the kernel function *κ*(*·, ·*) computes the relationship between pairs of channels. Formally, for an input time series **X** *∈* R*^c×t^*, composed of *c* channels and *t* time samples, the elements of the kernel matrices are defined as:

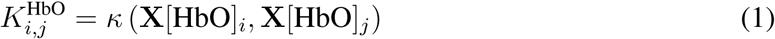

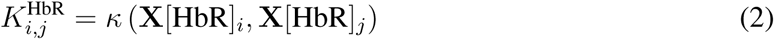

Here, **X**[HbO]*_i_* and **X**[HbR]*_i_* denote the *i*-th channel time series for HbO and HbR, respectively. The matrices **K**_HbO_ and **K**_HbR_ capture the intrinsic channel-to-channel relationships within their chromophore.

Instead of classifying HbO and HbR kernel matrices separately with different classifiers, one can combine them in a block diagonal matrix. A block diagonal matrix is a square matrix composed of smaller square matrices, or ”blocks,” along its diagonal and zero entries elsewhere. Therefore, our block diagonal matrix of **K**_HbO_ and **K**_HbR_ has the form:

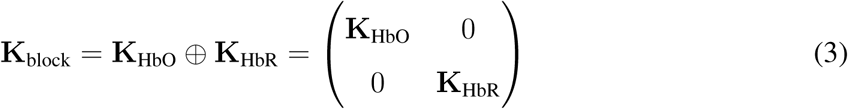

It is important to note that during hyperparameter tuning, we allow *κ*(*·, ·*) to be (non-linear) kernel functions as well as covariances with the different estimators, as described above. *κ*(*·, ·*) can also be different for HbO and HbR. To ensure the numerical stability of the kernel matrices, we additionally regularized **K**_HbO_ and **K**_HbR_ separately before combining them to **K**_block_. This regularization was done via shrinkage of the matrices, allowing different shrinkage parameters for each chromophore. This approach emphasizes flexibility for each oxygenation type and allows the classifier to be tuned separately to HbO and HbR data while ensuring the SPD property of the matrices. We termed this kernel matrix estimation “super kernel”.

In contrast, a full kernel matrix estimation does not permit such tailored optimization, which can limit performance. For full kernel matrix estimation, the approach is essentially the same as described earlier, however, only one kernel function is picked for both HbO and HbR. These full kernel matrices also include the interactions between HbO and HbR channels, while these interactions are set to zero for the block diagonal matrices. This means that the full kernel matrices can contain more information than the block diagonal matrices. However, they also lack the ability to tailor kernels separately for HbO and HbR.

#### 2.2.4 Co-activation comparison and mean kernel matrices

To test whether HbO and HbR show different co-activation patterns, we compared distances between the mean kernel matrices. For this, we first computed the mean kernel matrix for each task and for each participant, separately for HbO and HbR. Note that all mean kernel matrices were estimated with the Riemannian metric because it accounts for the non-Euclidean geometry of SPD kernel matrices, ensuring that the computed means and distances preserve the intrinsic structure of the data.^26, 27^ Next, we computed the pairwise Riemannian distances^28^ between all pairs of tasks (n = 28). The resulting distance vectors for HbO and HbR were transformed into discrete probability distributions. Subsequently, we computed the Wasserstein distance between these vectors. The Wasserstein distance, also known as Earth mover’s distance, can be thought of as the minimum ”cost” required to transform one distribution into the other. The ”cost” here is calculated based on the amount of probability mass that needs to be moved, and the distance it needs to be moved. Intuitively it represents a difference in shape between the two distributions. The logic is as follows: If HbO and HbR kernel matrices both contain the same information, the distribution of pairwise distances between the individual task pairs is similar. This would lead to a small Wasserstein distance. Likewise, if the two chromophores contain different information, and can hence differently differentiate between task pairs, the distance will be larger. We calculated the Wasserstein distance between HbO and HbR pairwise task distances for all participants and tested whether this distribution is different from zero with a one-sample t-test.

#### 2.2.5 Classification approach

The dataset used in this study consisted of eight different mental tasks with twelve trials each, for a total of 96 trials per participant. Given the relatively small size of the dataset, we opted not to split the data into dedicated training, validation, and test sets. Instead, we employed a repeated and stratified k-fold cross-validation approach, which is a common approach in machine learning scenarios with small datasets. Specifically, we applied 5-fold cross-validation with five repeats to compute the accuracy score for model comparisons. While this method is widely accepted and helps mitigate the variability in small datasets, it is important to note that it can overestimate the accuracy compared to using a separate test set. However, we performed the same approach for both traditional models as well as the Riemannian models. All computations were performed on an HPC cluster using ACME, scikit-learn, and pyriemann.^24, 25, 29^

#### 2.2.6 Relative improvement score

We calculated a relative improvement score for Riemann models compared to traditional models for each participant. This score normalizes the improvement by considering the proximity of the baseline accuracy to the maximum possible accuracy (i.e., 1). The relative improvement score is defined as follows:

Let *A*_baseline_ be the accuracy of the baseline model, and *A*_new_ be the accuracy of the new model.

The relative improvement score *RIS* is calculated using the formula:

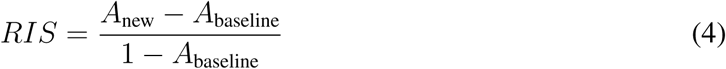

This formula scales the observed improvement by the remaining potential for improvement, given the baseline performance. The numerator, *A*_new_ *− A*_baseline_, represents the absolute improvement in accuracy achieved by the new model. The denominator, 1 *− A*_baseline_, represents the maximum possible improvement from the baseline model’s accuracy to perfect accuracy. By using this relative improvement score, we account for the fact that achieving improvements is more challenging as the baseline accuracy approaches 1.

#### 2.2.7 Statistical tests for model comparison

We compared the classification accuracies of traditional methods to the Riemann methods using paired *t*-tests. Since we used eight Riemann models and four traditional models, we determined the average accuracy for each participant and each model type using a bootstrap procedure (n = 1000). We determined the mean of the bootstrapped accuracy distribution and then performed the *t*-tests on these bootstrapped means.

For the best-performing model for the 2-choice and 8-choice comparison, we determined whether the repeated stratified 5-fold cross-validation accuracy was above chance level. The theoretical chance levels for the 2- and 8-choice comparisons are 50% and 12.5%, respectively.

However, these numbers are only valid for an infinite amount of samples. To better estimate classification performance, we used a permutation-based approach where we shuffled the labels (n = 500) and retrained models on the shuffled labels.^30^ All analyses used a significance threshold of *P <* 0.05 unless stated otherwise.

## 3 Results

### 3.1 Leveraging the inherent duality of fNIRS data with block diagonal matrices

To optimally use the complementary information that HbO and HbR can provide, we designed models that use block diagonal matrices for classification. This has the advantage that the manifold on which the data are classified contains both information on HbO and HbR that the model can leverage. The pipeline for fitting and estimating these models is illustrated in Fig 2.

**Fig 2.**
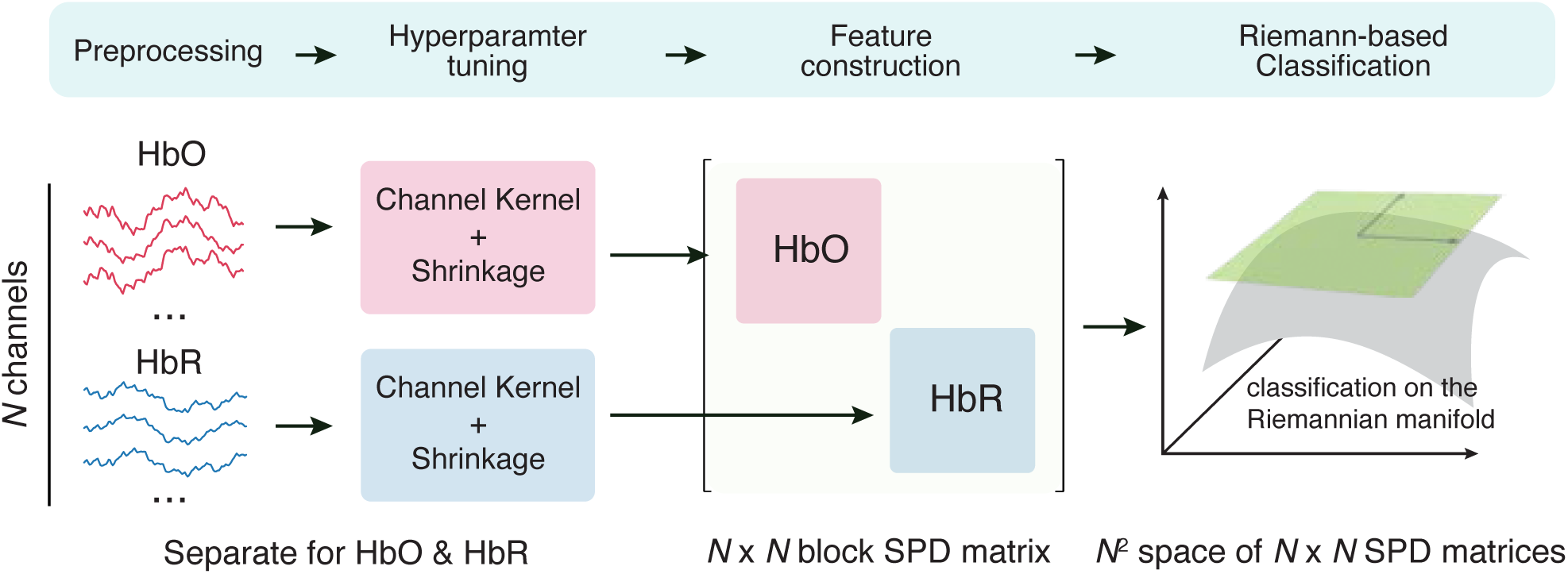
Processing pipeline of Riemannian fNIRS classifiers using block diagonal matrices as features. The data are first preprocessed according to standard pipelines. The subsequent hyperparameter tuning finds the best kernel functions and regularization for HbO and HbR separately. Each combination of hyperparameters is evaluated using a 5-fold stratified and repeated cross-validation (five repetitions). The resulting individual kernel matrices are then combined in a block diagonal matrix which serves as a feature for each trial. Several Riemannian-based classifiers were tested for classification of the fNIRS data of each participant.

The process begins with standard fNIRS preprocessing steps (detailed in the Methods), during which the data are separated into HbO and HbR signals. After preprocessing, the HbO and HbR data are processed separately (Fig 2). For each chromophore, we compute a kernel matrix with different estimators according to the hyperparameter set (Supplementary Table 3). This separation allows the hyperparameter search to identify the optimal function for each chromophore, enhancing the model’s sensitivity and performance. Following the estimation, the HbO and HbR matrices are regularized separately using shrinkage. The regularization step is crucial for stabilizing the matrices, especially given the high dimensionality of the data. Once regularized, the two matrices are combined into a block diagonal matrix, which is then used as input for a classifier (Fig 2).

This method offers a significant advantage by allowing the most efficient kernel and shrinkage parameters to be determined for each signal type individually.

We benchmarked twelve different classifiers in two different settings: Classifying eight tasks and classifying two tasks. For each setting, we evaluated four traditional machine learning models and eight Riemannian geometry-based models. The traditional models were LDA, LR, RF, and SVC. These models were trained using a traditional feature set comprising channel mean, variance, kurtosis, zero-crossings, and skewness.^22^ These features were computed per channel separately for HbO and HbR.

The eight Riemannian models are differentiated by variations in feature estimation and classifiers. We assessed a Riemannian-adapted SVC and LR in the tangent space of the Riemannian manifold. A full table of their hyperparameters can be found in Supplementary Table 3. The feature estimation techniques explored were as follows:

a. Full kernel matrix estimation: This approach concatenated HbO and HbR signals into a single 124 x 124 kernel matrix.
b. Block diagonal covariance matrix: This matrix retained the same dimensions as the simple covariance matrix but with off-diagonal blocks set to zero.
c. Block-kernel matrix: Here, the HbO and HbR blocks represented channel kernel functions instead of covariances.
d. Super-kernel block diagonal matrix: In this approach, the HbO and HbR blocks could be any kernel function, including covariances.

### 3.2 Hyperparameter tuning allows for individualization

Only a few studies have utilized channel relationships, such as kernel functions, to classify fNIRS data. However, those who have explored this approach demonstrated that kernels outperformed traditional fNIRS features for classifying mental tasks.^31^ For kernel matrices to be an effective feature for classification, it is crucial to record the activity from different brain regions as separated signals (minimizing mixture) and to obtain as many independent brain-activation measurements as possible. This can be optimally achieved by covering a large cortical area. Thus, our montage included 62 optode channels (Fig 3a), spanning the whole left hemisphere from occipital to frontal regions as well as parts of the right hemisphere. We sorted the individual channels of the kernel matrices according to their locations in the 10-20 EEG system in (anterior)-frontal, (front)-central, (centro)-parietal, occipital-parietal, temporal, and occipital. Fig 3a shows this sorting for two example kernel matrices (HbO and HbR) and the tasks “Mental Talking” and “Own Paradigm” (all eight tasks for this participant can be found in Supplementary Fig 2). Generally, for the group, kernel values were higher for HbO than HbR (*t* = 5.08*, P* = 2.26 *×* 10*^−^*^3^), likely due to the different signal-to-noise ratios of the two signals. While scaling covariance matrices affects the eigenvalues, it does not affect the eigenvectors. We tested whether HbO and HbR displayed different co-activation patterns by comparing the distributions of pairwise distances between all possible combinations of two tasks. We found that the average distance differs from zero, indicating that HbO and HbR pairwise distances contain different information structures (*t* = 4.68, *P* = 0.003). Generally, HbO showed a higher average distance between task pairs compared to HbR (*t* = 3.25, *P* = 0.017).

**Fig 3.**
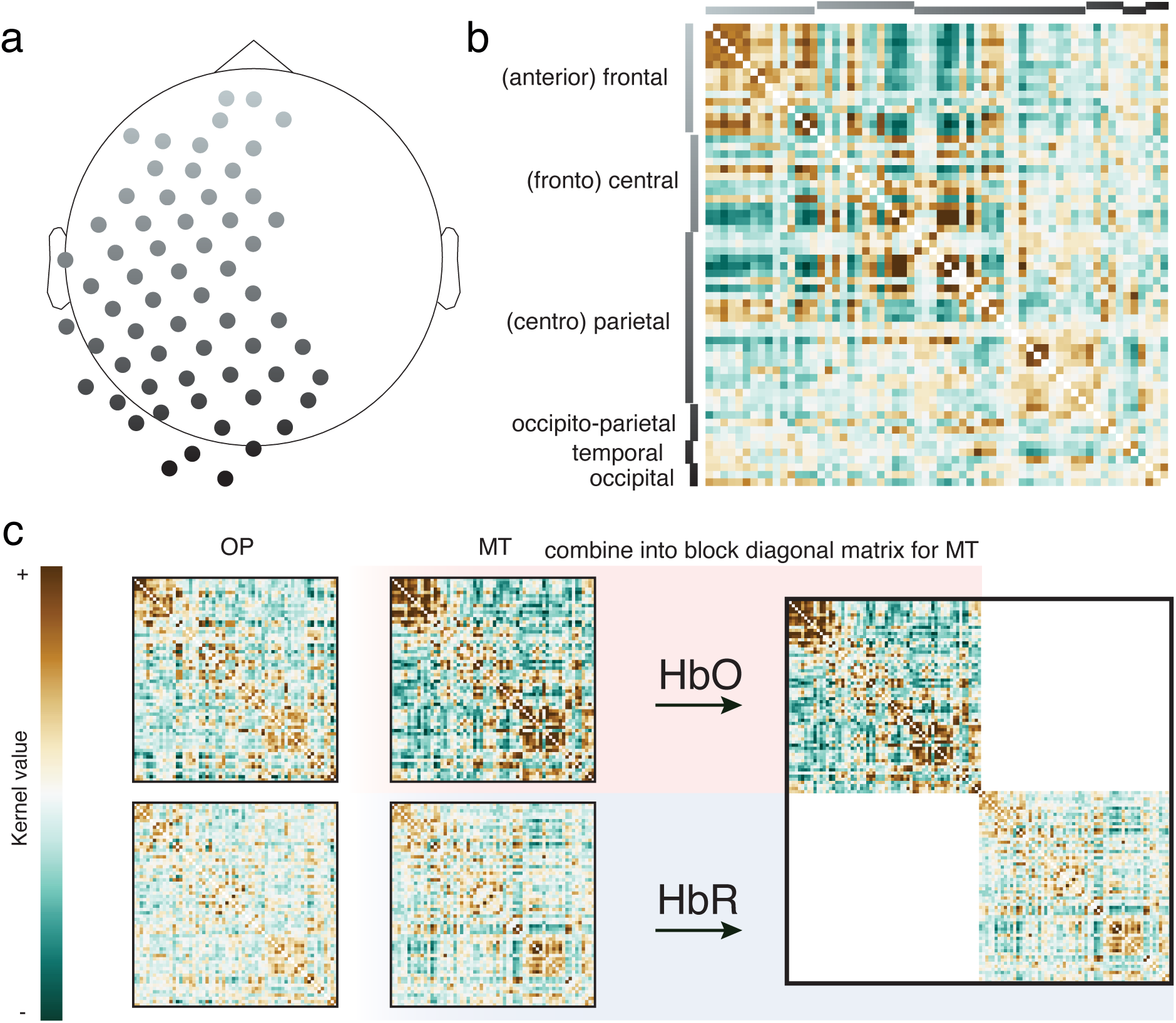
Example average kernel and block diagonal matrices. **a** FNIRS channel map. **b** Example kernel matrix for P01 and task MT. Rows and columns of the matrix are reordered according to the respective EEG locations of the channels. **c** Example average kernel matrices from participant P01 for HbO (top row) and HbR (bottom row) for task OP (left column) and task MT (right column). For task MT, the HbO and HbR kernel matrices are stacked into a block diagonal matrix on the right; the same is done for task OP but not illustrated here for space reasons. Note that the diagonal is set to zero for illustration purposes. MT: mental talking, OP: own paradigm.

The duality of fNIRS data suggests that one could use HbO and HbR as separate features for classification. However, adding more features to classification analyses does not necessarily improve classification accuracy.^32^ Riemannian geometry provides an elegant framework for combining different kernel matrices (features) into a single matrix by stacking them along the diagonal, resulting in a block diagonal matrix (Fig 3c). Thus, Riemannian geometry is particularly well-suited to manage the duality inherent in fNIRS data.

### 3.3 Riemannian models consistently outperform other models

As the final step of our model fitting process, we classify the previously estimated kernel matrices using either Riemannian-adapted or traditional classification models. Given that our dataset includes eight mental imagery tasks, it provides a valuable opportunity to test the potential of co-activation-type classification in distinguishing among multiple tasks. For this 8-choice classification, the traditional models (LDA, LR, RF, SVC) yielded a mean classification accuracy of 42% (all *P <* 0.01, permutation tests). In contrast, the Riemannian models demonstrated a higher mean accuracy of 62% (bootstrapped paired *t*-test, *t* = 7.70*, P* = 2.50 *×* 10*^−^*^4^; see Fig 4a).

**Fig 4.**
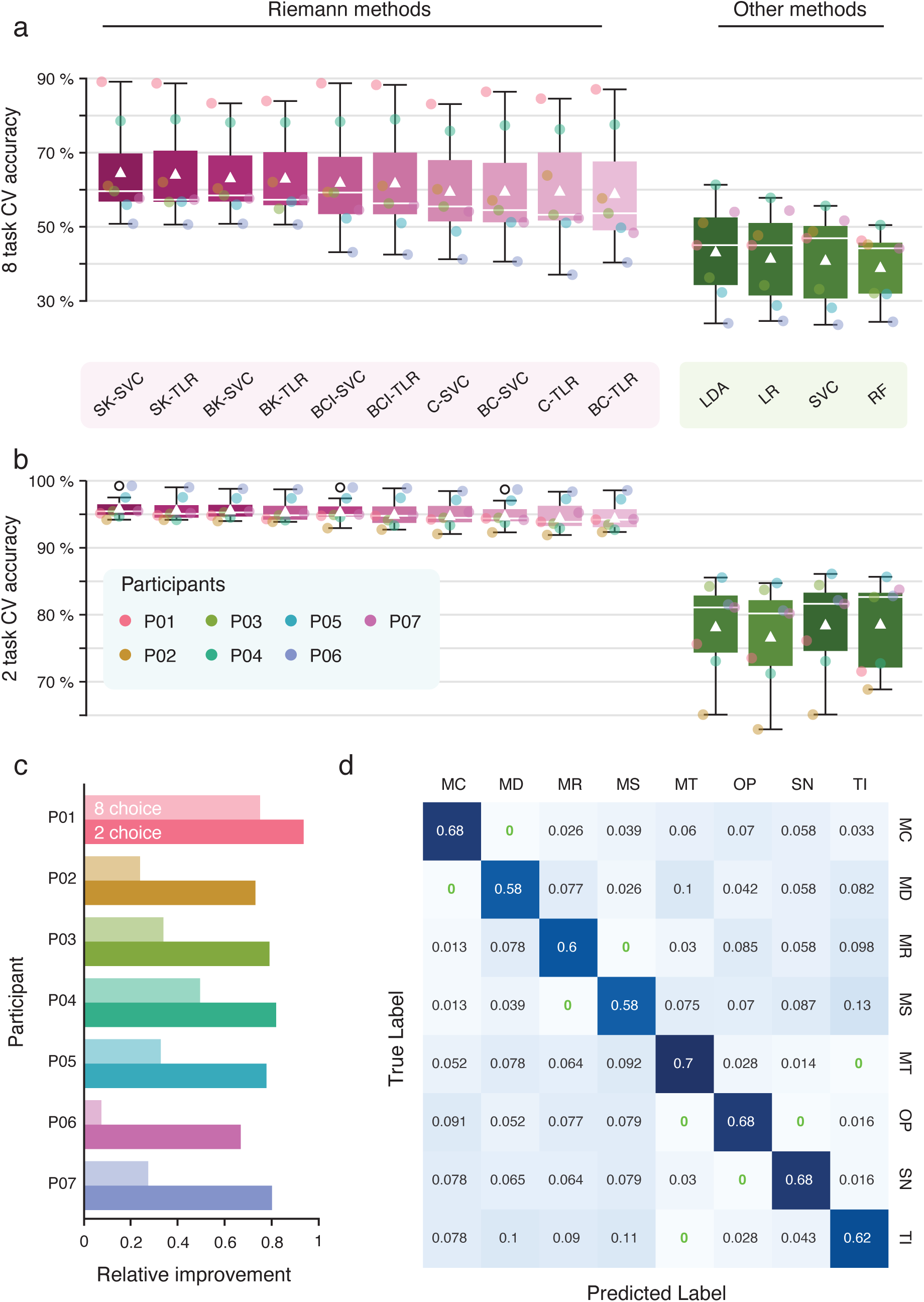
Classification results. Classification results of ten different Riemann-based methods (pink) and four traditional methods (green). **a** the mean accuracies for the one multi-class classification of all combinations of eight tasks. The box plot shows the median for each model as a white bar and the mean as a white triangle. Outliers are depicted as circles. **b** same as **a**, but for the 28 binary classifications of all combinations of two tasks (n = 7 participants in both cases). **c** Relative improvements of average accuracies from traditional to Riemann methods per participant. Lighter colors indicate relative improvements for the 8-choice paradigm and darker colors indicate relative improvements for all possible 2-choice combinations. **d** Mean confusion matrix for the 8-choice comparison, using the best-performing model (SK-SVC). Cells for tasks that were never confused with other tasks are marked in green. Abbreviations: SK: super-kernel, BK: block-kernel, C: covariance, BCI: block covariance individual, SVC: (Riemannian) support vector classifier, TLR: tangent space logistic regression, SVC: support vector classifier, RF: random forest, LDA: linear discriminant analysis, LR: logistic regression

In addition to evaluating the 8-choice classification performance across all tasks, we also explored a binary classification scenario by examining all possible pairwise combinations of the eight tasks, resulting in 28 different binary comparisons. Here, the traditional models achieved a mean accuracy of 78% with all classification accuracies above chance level (all *P <* 0.01, permutation tests). The Riemannian models outperformed the traditional models, achieving an average classification accuracy of 95% (bootstrapped paired *t*-test, *t* = 4.78*, P* = 3.05 *×* 10*^−^*^3^; see Fig 4b).

The “super-kernel SVC” model performed best in both the 8-choice and 2-choice comparisons (65% and 96% accuracy, respectively). Overall, models utilizing block diagonal matrices (n = 8) as features outperformed those based on full-kernel matrices (models: C-SVC, C-TLR) (n = 2) for both 2-choice comparison (*t* = 7.57*, P* = 2.77 *×* 10*^−^*^4^) and 8-choice comparison (*t* = 6.34*, P* = 7.23 *×* 10*^−^*^4^).

To quantify the improvement, we calculated a relative improvement score comparing the performance of traditional models to Riemannian models. This analysis revealed that the improvement was more pronounced in the 2-choice comparisons than in the 8-choice comparisons for all participants (*t* = *−*8.37*, P* = 1.58 *×* 10*^−^*^4^; see Fig 4c). Using our best-performing model, we analyzed the average confusion matrix for the 8-choice classification (Fig 4d). The results revealed that ”Mental Talking” (MT) had the highest accuracy in predicted versus true labels, with 70% correctly classified. This was followed closely by ”Mental Calculation” (68%), ”Own Paradigm” (68%), ”Spatial-Navigation Imagery” (68%), ”Tennis Imagery” (62%), ”Mental Signing” (58%), and ”Mental Drawing” (58%). Interestingly, some tasks were never confused with others. For instance, ”Mental Talking,” the most distinguishable task, was never mistaken for ”Tennis Imagery”. Similarly, ”Mental Calculation” was never confused with ”Mental Drawing”, ”Mental Drawing” was never confused with ”Mental Calculation”, ”Mental Rotation” was never confused with ”Mental Signing”, ”Mental Signing” was never confused with ”Mental Rotation”, and ”Own Paradigm” was never confused with ”Spatial-Navigation Imagery” or ”Mental Talking”, ”Spatial-Navigation Imagery” was never confused with ”Own Paradigm”, and ”Tennis Imagery” was never confused with ”Mental Talking”. These results suggest that certain task pairs may generate more orthogonal co-activation patterns than others, which can be effectively captured by our proposed methods. However, the classification accuracy for each task did not show a significant correlation with participants’ ratings of ease and pleasantness (all *P >* 0.05, Spearman rank correlation; see Supplementary Fig 1).

## 4 Discussion

In this study, we demonstrate that classification models based on Riemannian Geometry offer a powerful and robust method for classifying brain states with fNIRS. We show that Riemannian models consistently outperformed traditional models in both binary- and multi-choice classification tasks (see Fig 4). Additionally, our study underscores the benefits of leveraging the complementary information inherent to HbO and HbR signals captured by fNIRS. Stacking HbO and HbR signals in block diagonal matrices rather than full covariance matrices improves classification performance. Specifically, the combination of a block diagonal matrix as a feature together with a Riemannian Support Vector Classifier (SVC) achieved the highest accuracy among all models in both the 8-choice and 2-choice classification (see Fig 4), with an average of 65% and 96% cross-validation accuracy, respectively.

### 4.1 The Benefits of Combining Riemannian Geometry with a Block Diagonal Approach

Riemannian models consistently outperformed traditional models such as LR, SVC, LDA, and RF (see Fig 4). This difference in performance likely stems from the distinct features each approach utilizes for classification. While traditional models rely on statistical features computed from individual channels,^12, 13, 22^ Riemannian models harness information from channel variances and interactions.^15^ Given that brain processing is coordinated by network activity, and is therefore based on interactions, (non)-linear channel interactions are inherent.^33^ Unlike other statistical features like mean, kurtosis, or zero-crossings, kernel matrices which serve as the central feature in Riemannian approaches, can capture these interactions. For these kernel matrices, non-linear channel kernels can offer an improvement over regular covariances.^23^

Among the Riemannian models, those employing block diagonal matrices as features outperformed models using full kernel matrices at the group level (see Fig 4). The block diagonal approach allows for the concatenation of multiple SPD matrices, each estimated using potentially different kernel functions and regularization strengths. These models have a larger hyperparameter set, and this additional complexity allows these models to be more finely tailored to the specific characteristics of the two chromophores. The increased flexibility in hyperparameter tuning likely contributed to the superior performance compared to full kernel models. Given that in our data there appears to be minimal meaningful interaction between the HbO and HbR signals, estimating a full kernel matrix may unnecessarily increase the dimensionality of the manifold without further improving classification accuracies. By setting the interactions between HbO and HbR to zero, block diagonal matrices eliminate these redundancies, resulting in a manifold with lower dimensionality. This submanifold, corresponding to the block diagonal matrix, is therefore more compact and likely better suited to capture the essential structure of the data.

The best-performing model in our study was a block diagonal Riemannian SVC (see Fig 4). As a powerful linear classifier, the SVC is known for its robust generalization capabilities. When data are not linearly separable in their native space, SVC can map them to a higher-dimensional space where linear separation becomes possible, making it highly efficient in capturing potential non-linear interactions.^34^ Although SVCs traditionally operate with Euclidean feature vectors, they can be extended to incorporate a Riemannian kernel, allowing for kernel matrices to be used as input features.^35^

In this study, we compared a regular SVC with traditional features to the Riemannian SVC with (block) kernel matrices as features. The Riemannian SVC, based on the block diagonal approach with separate HbO and HbR tuning, stands out as the most effective method for classifying fNIRS mental imagery data in our study.

### 4.2 Integration of Riemannian Geometry in fNIRS-based Applications

A rapidly growing field of fNIRS applications is brain-computer interfacing for motor-independent communication and control, where voluntarily generated brain activations are used to encode computer commands. In these systems, the ability to accurately classify distinct patterns of brain activity is crucial for enabling effective communication and control. As fNIRS-based brain-computer interfaces (BCIs) continue to evolve,^36^ integrating advanced analytical techniques like Riemannian Geometry has the potential to significantly enhance the precision and reliability of these systems. Similarly, Riemannian Geometry has successfully been used to advance EEG-based BCIs^35, 37–40^ and extending this approach to fNIRS represents a natural progression. By using mental-imagery tasks, crucial for BCI-applications, we show that applying Riemannian Geometry to fNIRS data has great potential for the BCI field. Most prominent is the significant increase in classification accuracy that can be achieved with Riemannian models compared to traditional models (see Fig 4a). Additionally, Riemannian models are computationally efficient and require comparatively less training data than other modern models like neural networks. Lastly, tools like the pyrie-mann package simplify the implementation of Riemannian models, making them accessible to researchers across various disciplines. These advantages could make fNIRS a viable alternative to other BCI methods, like fMRI and EEG.

Currently, EEG remains the predominant method in real-world BCI applications,^41^ with several established classification methods based on Riemannian Geometry.^16, 42–45^ Our study demonstrates that Riemannian Geometry may also be well-suited for fNIRS-based BCIs. Thus, the combination of EEG and fNIRS in hybrid BCIs represents a promising future direction.^46^ Future studies could identify co-activation patterns between these two modalities, potentially enhancing BCI performance by leveraging the complementary information from both techniques within a unified Riemannian Geometry framework. The integration of EEG with fNIRS might proceed according to the integration of HbO and HbR exemplified here, namely using block diagonal matrices. This integration could further significantly advance BCI technology, offering more robust and versatile systems in which the unique advantages of fNIRS and EEG can optimally be exploited. For practical BCI applications, it may be advantageous to focus on a smaller subset of tasks, selecting those that are most orthogonal to each other to enhance classification accuracy. Our dataset, which includes eight mental imagery tasks, offers a unique opportunity to identify which tasks are most distinguishable using Riemannian classification (see 4). We identified ”Mental Talking” as the most easily distinguishable task using our analysis approach, likely due to its intuitive nature and close relation to natural thought processes. Interestingly, participant ratings supported this, with “Mental Talking” ranking second highest for ”easiness” and third for ”pleasantness” (see Supplementary Fig 1). Next to the mentioned BCI applications, fNIRS-based classifications us-ing Riemannian Geometry could also have significant implications for other clinical applications considering its accuracy, computational efficiency, and accessibility. As an example, ongoing research in our lab applies the novel method to crucially advance earlier attempts for the brain-based diagnosis of remaining awareness in patients with disorders of consciousness.^47^

### 4.3 Limitations and Future Directions

Despite the promising results, several limitations must be considered. The study involved only seven participants, which limits the generalizability of the findings. However, it is important to note that clinical (e.g., BCI) research often prioritizes single-subject analyses, where generalization to the population is less important or even not necessary. Note also, that despite the limited sample size and the correspondingly low sensitivity, our findings consistently demonstrated that Riemannian models significantly outperformed traditional models at the group level.

Another limitation is the absence of a dedicated train-test split, which might lead to an over-estimation of classification accuracy. Rather, we use cross-validation accuracy, which can inflate real-world performance. This could in principle be mitigated by a train-test split of the available data, which was however not feasible due to the small number of trials per subject. However, this does not compromise the validity of our comparative results, as the cross-validation approach was used consistently across all tested models, ensuring no systematic bias or interaction. Additionally, Riemannian models required more extensive hyperparameter tuning than traditional models. Yet, this extended tuning process allowed us to optimize model performance and reach superior performance. Enhanced model sensitivity enables the detection of subtle neural effects, making this approach valuable for advancing our understanding of brain function. While accuracy is similarly important for BCI applications, time is often a limiting factor in applied fNIRS-BCI settings. BCIs using fNIRS generally work on longer time scales due to the characteristic time delay of the hemodynamic response. Additionally, conducting a large hyperparameter tuning might lead to long fitting times. Thus, it may be necessary to reduce the parameter space in BCI and other clinical contexts to achieve an optimal balance between model-fitting time and accuracy. Alternatively, a Riemannian model with a single kernel matrix and a subset of channels could offer a solution that requires less powerful hardware while still surpassing traditional approaches.

In the current study, ”Mental Talking” was the most distinguishable task out of all eight tasks. Of note is that mental talking may have elicited unconscious movements of the tongue or yaw. These subliminal movements could have contributed to the classification by increasing motor cortical activation. However, previous studies have used “Mental Talking” successfully for fNIRS-based classification in a movement-disabled patient,^48^ further supporting the distinguishability of the task. Future studies may measure electromyography to detect potentially confounding muscle activity. Note that any such subliminal movements would have been equally available to all classification methods and cannot explain the observed differences between methods.

Future research might investigate the integration of Riemannian Geometry with deep-learning techniques to further improve classification performance, a strategy that has shown promise in EEG.^49^ To validate the utility of both existing and novel models, they should be tested across diverse settings, from basic research to applied contexts, to determine which models are most effective in various scenarios.

## Acknowledgments

The authors gratefully acknowledge the volunteers for their participation in this study, as well as the contributions of former students involved in pilot phases of this research, including Thomas Emmerling, Martin Sona, Lizette Heine, and Caroline Friedrich. This research was funded by the Netherlands Organization for Scientific Research (NWO; RUBICON 446-09-010 and Vidi-Grant VI.Vidi.191.210 to BS) and the European Community’s Seventh Framework Programme (FP7/2007-2013) under grant agreement PITN-GA-2011-29001. The funding sources had no involvement in the study’s design, data collection, analysis, or interpretation.

## Appendix A: Supplementary Figures

**Supplementary Fig 1.**
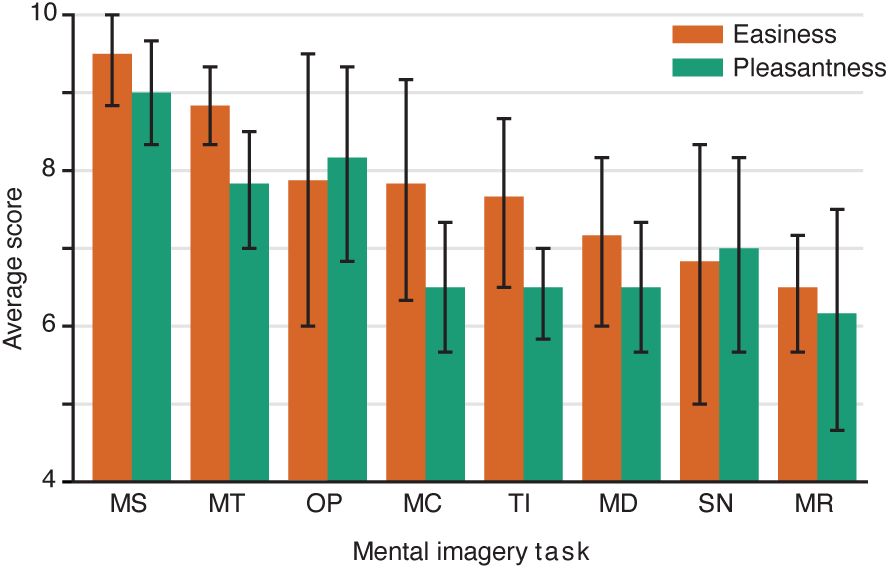
Task easiness and pleasantness ratings.

**Supplementary Fig 2.**
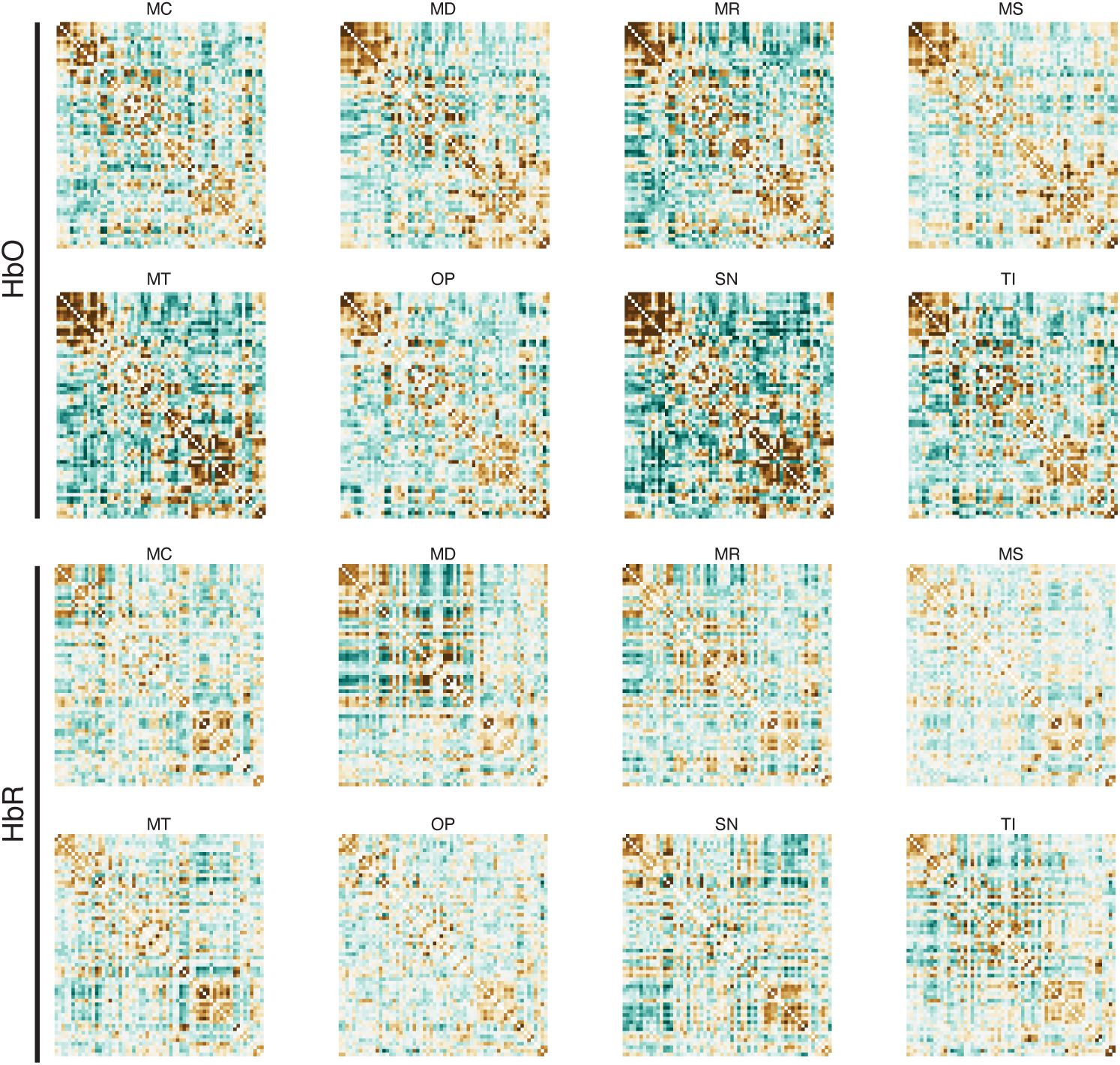
Example kernel matrices for participant P01 are shown for all eight tasks, with HbO matrices displayed in the top two rows and HbR matrices in the bottom two rows.

## Appendix B: Supplementary Tables

**Supplementary Table 1.**
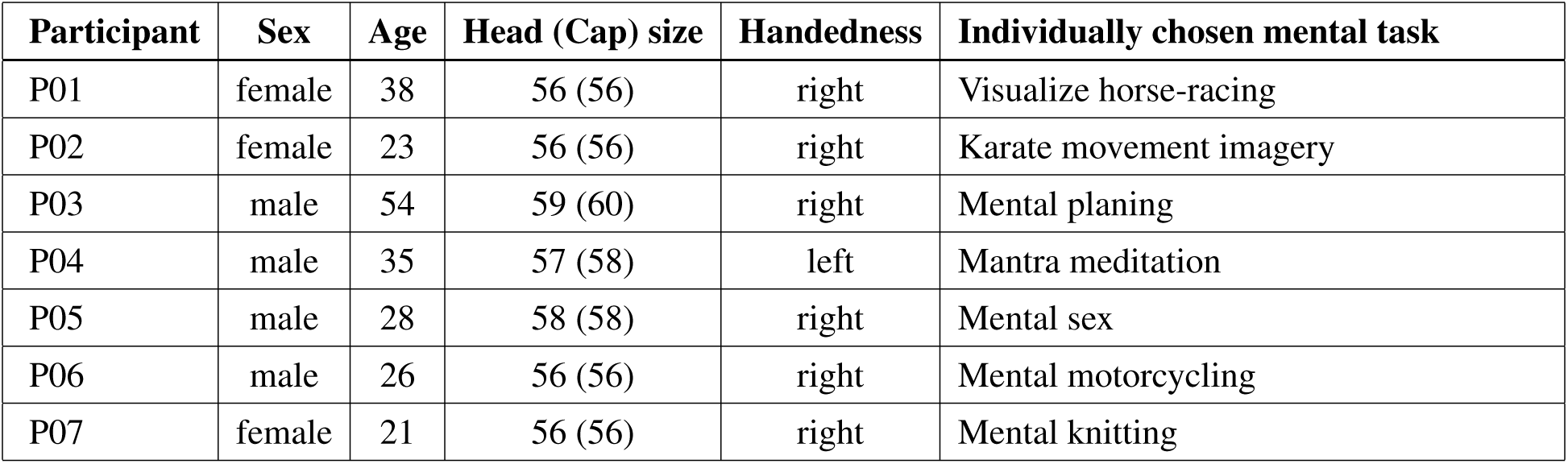
Participant Characteristics. The table presents the characteristics of each participant, including sex, age, head (cap) size, handedness, and individually chosen mental task.

**Supplementary Table 2.**
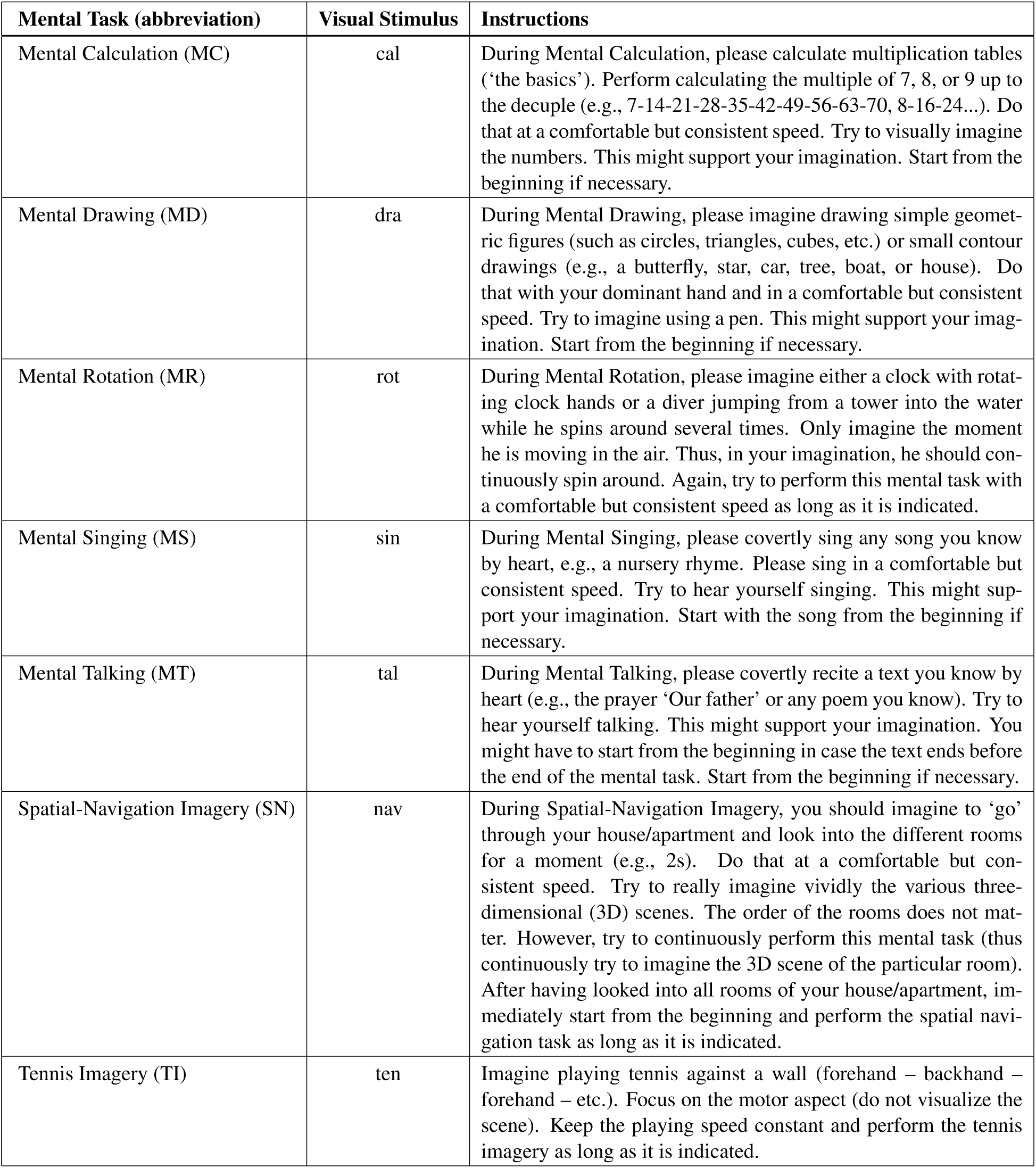
Overview of the mental tasks performed by participants in the current study, including abbreviations in brackets. The visual stimulus presented during the experimental session was limited to three characters (2nd column). Detailed instructions on how to perform each task were provided to participants before the experimental session.

| **Mental Task (abbreviation)** | **Visual Stimulus** | **Instructions** |
| --- | --- | --- |
| Mental Calculation (MC) | cal | During Mental Calculation, please calculate multiplication tables ('the basics'). Perform calculating the multiple of 7, 8, or 9 up to the decuple (e.g., 7-14-21-28-35-42-49-56-63-70, 8-16-24...). Do that at a comfortable but consistent speed. Try to visually imagine the numbers. This might support your imagination. Start from the beginning if necessary. |
| Mental Drawing (MD) | dra | During Mental Drawing, please imagine drawing simple geometric figures (such as circles, triangles, cubes, etc.) or small contour drawings (e.g., a butterfly, star, car, tree, boat, or house). Do that with your dominant hand and in a comfortable but consistent speed. Try to imagine using a pen. This might support your imagination. Start from the beginning if necessary. |
| Mental Rotation (MR) | rot | During Mental Rotation, please imagine either a clock with rotating clock hands or a diver jumping from a tower into the water while he spins around several times. Only imagine the moment he is moving in the air. Thus, in your imagination, he should continuously spin around. Again, try to perform this mental task with a comfortable but consistent speed as long as it is indicated. |
| Mental Singing (MS) | sin | During Mental Singing, please covertly sing any song you know by heart, e.g., a nursery rhyme. Please sing in a comfortable but consistent speed. Try to hear yourself singing. This might support your imagination. Start with the song from the beginning if necessary. |
| Mental Talking (MT) | tal | During Mental Talking, please covertly recite a text you know by heart (e.g., the prayer 'Our father' or any poem you know). Try to hear yourself talking. This might support your imagination. You might have to start from the beginning in case the text ends before the end of the mental task. Start from the beginning if necessary. |
| Spatial-Navigation Imagery (SN) | nav | During Spatial-Navigation Imagery, you should imagine to 'go' through your house/apartment and look into the different rooms for a moment (e.g., 2s). Do that at a comfortable but consistent speed. Try to really imagine vividly the various three-dimensional (3D) scenes. The order of the rooms does not matter. However, try to continuously perform this mental task (thus continuously try to imagine the 3D scene of the particular room). After having looked into all rooms of your house/apartment, immediately start from the beginning and perform the spatial navigation task as long as it is indicated. |
| Tennis Imagery (TI) | ten | Imagine playing tennis against a wall (forehand – backhand – forehand – etc.). Focus on the motor aspect (do not visualize the scene). Keep the playing speed constant and perform the tennis imagery as long as it is indicated. |

**Supplementary Table 3.** Overview of hyperparameters used during models fitting.

| **Hyperparameter Category** | **Values** |
| --- | --- |
| Kernel Metrics | [rbf, polynomial, laplacian, cosine, poly, corr, cov, scm, lwf, oas, hub, sch, tyl] |
| Matrix shrinkages | [0, 0.01, 0.1] |
| Riemann SVC C Values | [0.1, 1, 10, 100] |
| Riemann SVC Metrics | [euclid, riemann, logeuclid] |
| Tangent Space Logistic Regression C Values | [0.1, 1, 10, 100] |
| LDA Solver | [svd, lsqr, eigen] |
| LDA Shrinkage Values | 50 linearly spaced between 0 and 1 |
| Regular SVC C Values | [0.1, 1, 10, 100] |
| Regular SVC Kernels | [linear, poly, rbf, sigmoid] |
| Random Forest Estimators | [50, 100, 200] |
| Random Forest Max Depth | [None, 10, 20, 30] |
| Random Forest Min Samples Split | [2, 5, 10] |
| Random Forest Min Samples Leaf | [1, 2, 4] |
| Random Forest Max Features | [auto, sqrt, log2] |
| Logistic Regression C Values | [0.1, 1, 10, 100] |

